# Mechanistic Picture for Monomeric Human Fibroblast Growth Factor 1 Stabilization by Heparin Binding

**DOI:** 10.1101/2020.08.20.228056

**Authors:** Vivek Govind Kumar, Shilpi Agrawal, Thallapuranam Krishnaswamy Suresh Kumar, Mahmoud Moradi

## Abstract

Human fibroblast growth factor (FGF) 1 or hFGF1 is a member of the FGF family that is involved in various vital processes such as cell proliferation, cell differentiation, angiogenesis and wound healing. hFGF1, which is associated with low stability *in vivo*, is known to be stabilized by binding heparin sulfate, a glycosaminoglycan that aids the protein in the activation of its cell surface receptor. The poor thermal and proteolytic stability of hFGF1 and the stabilizing role of heparin have long been observed experimentally; however, the mechanistic details of these phenomena are not well understood. Here, we have used a combination of microsecond-level equilibrium molecular dynamics (MD) simulations, and state-of-the-art enhanced sampling MD simulations to quantitatively characterize the structural dynamics of monomeric hFGF1 in the presence and absence of heparin hexasaccharide. We have observed a conformational change in the heparin-binding pocket of hFGF1 that occurs only in the absence of heparin. Several intramolecular hydrogen bonds were also identified within the heparin-binding pocket, that form only when hFGF1 interacts with heparin. The loss of both intermolecular and intramolecular electrostatic interactions in the absence of heparin plausibly leads to the observed conformational change. This conformational transition results in increased flexibility of the heparin-binding pocket and provides an explanation for the susceptibility of *apo* hFGF1 to proteolytic degradation and thermal instability. The hFGF1-heparin interaction has also been quantified using absolute binding free energy calculations. Binding affinity (K_d_) estimates determined computationally using our novel MD approach are in good quantitative agreement with experimental K_d_ values from isothermal titration calorimetry experiments. The successful application of a combination of microsecond-level MD and accurate free energy calculations to explain the heparin-mediated stabilization of hFGF1 at a quantitative level, represents a promising approach for studying complex biomolecular interactions between proteins and their binding partners at a detailed molecular level using rigorous physics-based simulation techniques.

## INTRODUCTION

Thanks to the ever-increasing power of computers, improved force fields, and high-throughput modeling, all-atom MD is now routinely used to simulate proteins in simplified but explicit aqueous/membrane environments. MD simulations combine the high spatial resolution of experimental methods such as X-ray crystallography with the high temporal resolution of experimental methods such as single-molecule FRET spectroscopy^1,2^. However, many MD studies implicitly assume that local conformational transitions observed in short, nanosecond-level simulations can be used to describe global protein conformational transitions that typically occur on microsecond or millisecond time scales^3,4^. We have recently demonstrated that longer microsecond-level simulations are essential for a more precise statistical characterization of both local and global conformational transitions^5^. While microsecond-level MD simulations provide a more accurate description of protein structural dynamics as compared to shorter simulations, efficient sampling of the conformational landscape remains a major problem and requires access to longer timescales.

Accurate study of chemo-mechanical coupling in proteins requires the use of microsecond-level MD simulations in conjunction with enhanced sampling techniques and free energy calculations that go beyond microseconds. An effective computational protocol for characterizing functionally important conformational changes, using a combination of several enhanced sampling techniques without compromising the chemical details, has recently been developed^6,7,8,9^. Extensive structural information obtained using this strategy would allow for realistic experimental validation of the computational results. Here, we use a similar strategy to investigate the conformational and structural dynamics of monomeric hFGF1 (PDB entry: 1RG8)^10^ and the chemo-mechanical coupling between hFGF1 and heparin hexasaccharide, its glycosaminoglycan (GAG) binding partner.

Fibroblast growth factors (FGFs) are signaling proteins that are involved in an extensive variety of physiological processes^11,12,13^. The biological activity of FGFs is regulated through interactions with linear anionic polysaccharides called glycosaminoglycans (GAGs), which facilitate binding to specific receptors on the cell surface (FGFRs)^14,15,16,17,18,19,20,21^. Human acidic fibroblast growth factor (hFGF1) is an important signaling molecule expressed in embryonic and adult tissues for angiogenesis, cell proliferation and differentiation, tumor growth, neurogenesis and wound healing^14,22^. Glycosaminoglycans (GAGs) consist of a class of negatively charged and large linear polysaccharides formed of repeating disaccharide units in which a uronic acid (either glucuronic acid or iduronic acid) moiety is combined with an amino sugar (either *N-acetyl*-D-glucosamine or *N-acetyl*-D-galactosamine)^23,24^. Heparin is a GAG made up of 2-O-sulfated iduronic acid and 6-O-sulfated, N-sulfated glucosamine (IdoA(2S)-GlcNS(6S)), connected by *α*-(1→4) glycosidic linkages^25^. The anionic nature of GAGs leads to electrostatic interactions with positively charged (Lysine/Arginine-rich) regions of their target proteins^23,24^. The hFGF1-heparin complex is the most broadly studied protein-GAG complex^26,27^.

The interaction of hFGF1 with specific heparin sulfate proteoglycans may be influenced by the flexibility of the heparin-binding pocket^28^. In addition to the structural features of hFGF1, GAG sulfation patterns also determine the functionality and specificity of protein-GAG interactions^29,30^. hFGF1 is known to selectively recognize the GlcNS-IdoA2S-GlcNS sulfation motif^31^. DiGabriele *et al.*^32^ crystallized a dimeric hFGF1-heparin sandwich complex (PDB entry: 2AXM) and showed that heparin binding does not result in any major conformational changes within hFGF1^32,33^. Solution nuclear magnetic resonance (NMR) and experimental binding studies suggest that a monomeric hFGF1-heparin complex is also fully functional^27,34^. *Apo* hFGF1 shows relatively low thermal stability and is known to be susceptible to thermal degradation^35,36^. Binding to heparin sulfate proteoglycans is thought to protect hFGF1 against proteolytic degradation^37,38^.

Our microsecond-level all-atom equilibrium MD simulations reveal that a conformational change occurs in the heparin-binding pocket of hFGF1 in the absence of heparin. We postulate that this conformational change is responsible for the susceptibility of unbound hFGF1 to thermal instability, as seen in equilibrium unfolding experiments. We have also studied the intermolecular interactions of the hFGF1-heparin complex and the intramolecular interactions that are unique to heparin-bound hFGF1 in order to obtain a clearer picture of the heparin-mediated stabilization. In addition to qualitative analysis of the complex, we have also quantified the hFGF1-heparin interaction in terms of its absolute binding free energy. Recent computational studies have attempted to calculate the binding free energy of the hFGF1-heparin complex using the MM-GBSA approach^39^. Adequate sampling of ligand movements with respect to the protein is essential for quantifying the configurational entropy arising from the hFGF1-heparin association^40,41,42^. Free-energy calculation methods such as MM-PBSA/GBSA typically ignore a portion of this configurational entropy^40,43^. Here, we have calculated the absolute binding free energy of the hFGF1-heparin complex using a combination of non-equilibrium steered molecular dynamics (SMD) simulations and bias-exchange umbrella sampling (BEUS)^6,44,45,46^. Our computationally determined binding affinity estimates are in good agreement with the results of the isothermal titration calorimetry (ITC) experiments.

## RESULTS AND DISCUSSION

The putative role of heparin is to prevent the degradation of hFGF1. However, the specifics of this heparin-mediated stabilization are still unclear. To address this, we have used a combination of experimental and computational methods to compare and characterize the *apo* and heparin-bound forms of hFGF1.

We have performed three unbiased all-atom MD simulations of monomeric hFGF1, each for 4.8 μs. One *apo* and two heparin-bound models were simulated in the presence of explicit water. The *apo* model and one of the heparin-bound models (Model 1) are based on the crystal structure of monomeric *apo* hFGF1 (PDB entry: 1RG8)^10^. The second heparin-bound model is extracted as a monomeric model from the crystal structure of dimeric heparin-bound hFGF1 (PDB entry: 2AXM)^32^ (Supplemental Figure S1). Heparin hexasaccharide from the latter (2AXM) structure was added to the former (1RG8) monomeric structure in order to build heparin-bound Model 1. Similarly, one of the monomers in the 2AXM structure was removed to build heparin-bound Model 2. The heparin-bound models are virtually identical and involve an hFGF1 monomer bound to heparin hexasaccharide. As the only difference between the *apo* and heparin-bound models is the presence or absence of heparin, we can make meaningful comparisons between all three sets of simulations.

### A conformational change occurs in the heparin-binding pocket of the apo model

A visual inspection of the trajectories clearly shows that the heparin-binding pocket (residues 126-142) of the *apo* model becomes elongated and extends further outward and away from the core beta-trefoil structure after approximately 2 μs (Figure 1A). This conformational change is not observed in either of the two heparin-bound models (Figure 1B, Supplemental Figure S2A). Comparing the internal root mean square deviation (RMSD) of the hFGF1 monomer from each system reveals that the presence of heparin hexasaccharide stabilizes the protein and prevents this conformational change from occurring (Figure 1C, Supplemental Figures S2A-B). All 3 models initially have internal RMSD values of approximately 1 Å. Both heparin-bound models settle down into a stable conformation within 0.5 μs (RMSD=1.5 to 2 Å approx.) (Figure 1C, Supplemental Figures S2A-B). On the other hand, the *apo* model clearly undergoes a conformational change after 2 μs (RMSD=3 Å approx.) (Figure 1C, 1A). This new conformation then remains stable for around 2.8 μs (Figure 1C, 1A).

**Figure 1.**
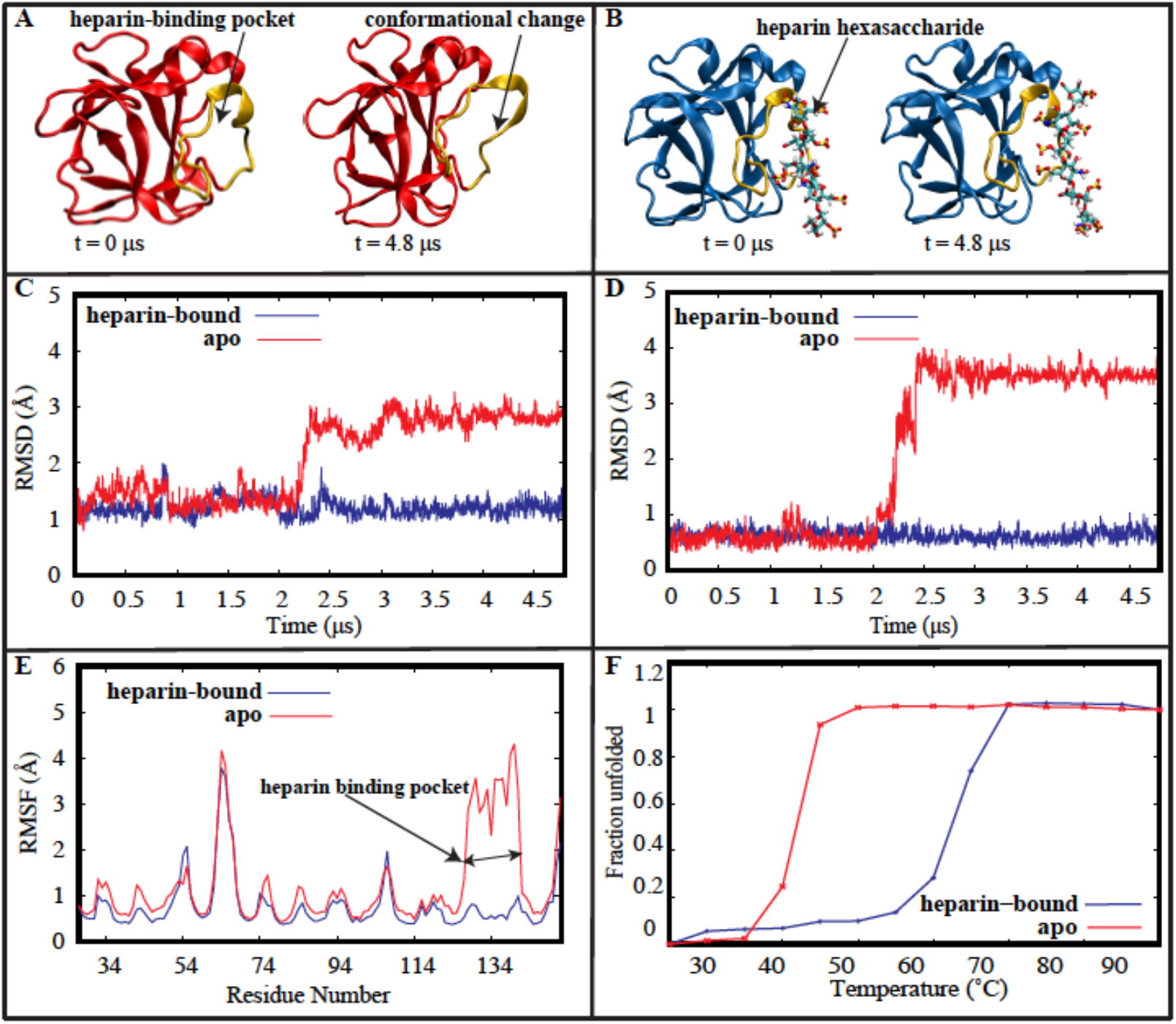
Conformational change in the heparin-binding pocket of *apo* hFGF1. (A,B) Cartoon representation of *apo* (red) and heparin-bound (blue) hFGF1 at the beginning and end of the 4.8-μs simulations. The heparin-binding pocket (gold) moves away from the beta-trefoil core of the *apo* protein. (C,D) Root mean square deviation (RMSD) time series for the *apo* (red) and heparin-bound (blue) models of hFGF1 protein (C) and its heparin binding pocket (D). (E) Root mean square fluctuation (RMSF) estimations for the *apo* (red) and heparin-bound (blue) models of hFGF1. (F) Thermal denaturation data for hFGF1 in the absence (red) and presence (blue) of heparin. The presence of heparin causes the T_m_ value to increase by around 20°C, indicating that heparin stabilizes the protein.

A comparison of the internal RMSD of the heparin-binding pocket reveals that this region plays a key role in the differential behavior of the *apo* (RMSD=4 Å approx.) and heparin-bound models (RMSD=0.5 Å approx.) (Figure 1D, Supplemental Figures S2C-D). This indicates that the absence of any interactions with heparin contributes to the decreased stability of the *apo* model. These results are also supported by the root mean square fluctuation (RMSF) data for each model, which was calculated for the C_α_ atoms of all protein residues (Figure 1E, Supplemental Figure S2E-F). All three models show similar trends in the fluctuations for different regions, with the exception of the heparin-binding pocket. As expected, the heparin-binding pocket is much more flexible in the *apo* model than in the heparin-bound models.

Thermal denaturation experiments were performed on monomeric hFGF1, in the absence and presence of heparin hexasaccharide, to further validate our computational results. The T_m_ value for the *apo* experimental model was approximately 42°C while the T_m_ value for the heparin-bound experimental model was approximately 62.5°C (Figure 1F). The presence of heparin thus increases the T_m_ value by around 20°C, indicating that heparin stabilizes the protein. These observations are in qualitative agreement with the computational RMSD/RMSF data.

### Unique salt-bridge interactions facilitate the conformational change in the apo model

The conformational change that occurs in the *apo* model is localized in the heparin-binding pocket. Electrostatic interactions between positively charged residues in the heparin-binding pocket and negatively charged residues in the beta-trefoil core help stabilize the new conformation. We have identified two salt bridge interactions that are unique to the *apo* model. They do not form in the two heparin-bound models (Figure 2A-D, Supplemental Figure S3A-D). D84 of the beta-trefoil core interacts with K132 of the heparin-binding pocket (Figure 2A-B), while D46 of the beta-trefoil core interacts with K127 of the heparin-binding pocket (Figure 2C-D). When the heparin-binding pocket starts moving during the conformational change, a weak salt bridge forms between D46 and K127 (Figure 2C-D). This is followed by the formation of a stronger salt bridge between D84 and K132 when the heparin-binding pocket becomes elongated and is extended outward (Figure 2A-B) and away from the beta-trefoil core. Both K127 and K132 are known to interact with negatively charged heparin residues^34^. Interactions with negatively charged residues of the beta-trefoil core possibly compensate for the absence of interactions with heparin. Together, these salt bridges play a key role in stabilizing the new conformation of the heparin-binding pocket for almost 2.8 μs.

**Figure 2.**
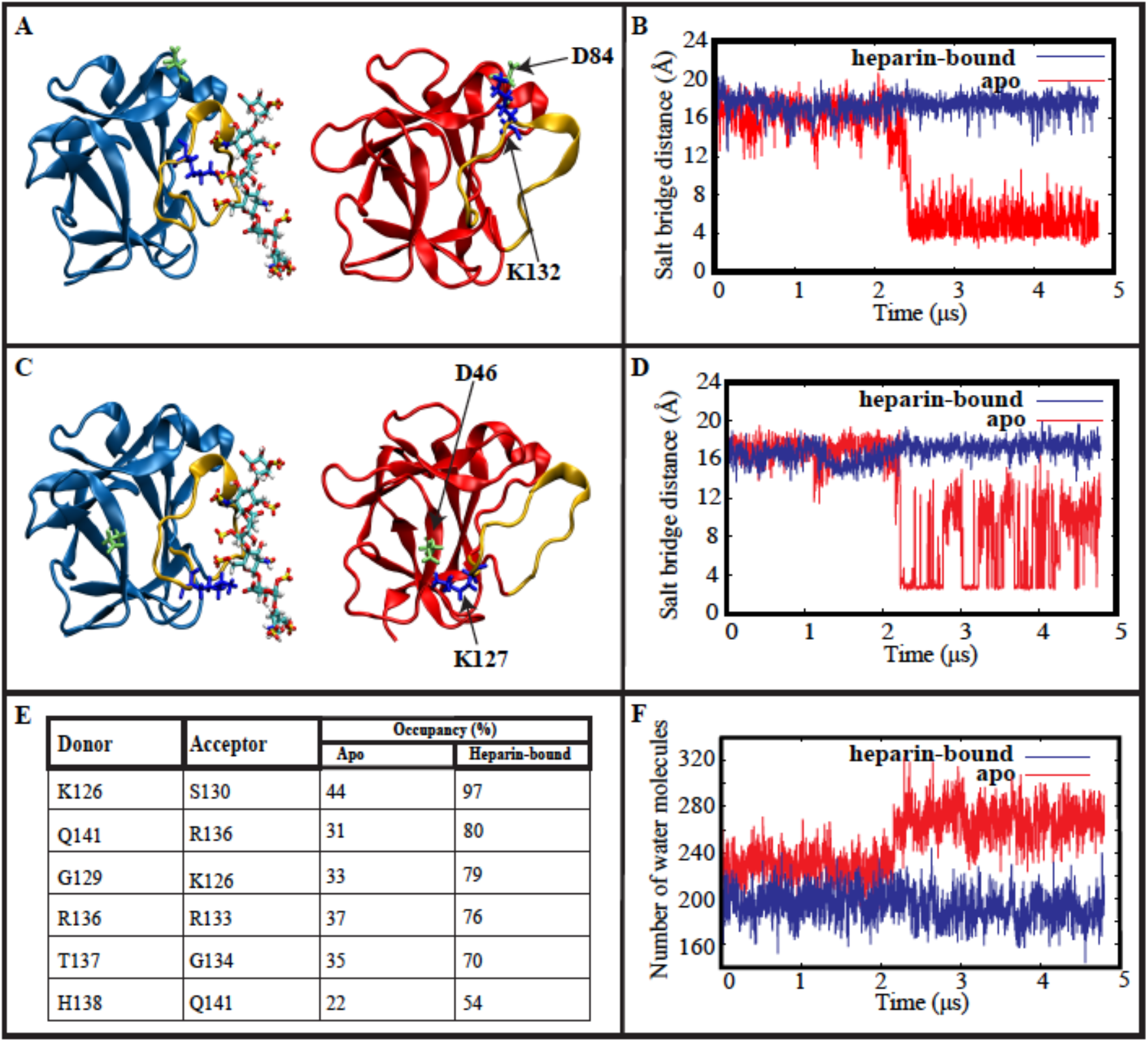
Unique salt-bridge interactions facilitate the conformational change in the *apo* model. (A) K132 (blue) of the heparin-binding pocket (gold) forms a salt-bridge with D84 of the beta-trefoil core in the *apo* model (red). This interaction does not form in the heparin-bound protein (blue). (B) Time series of the D84-K132 donor-acceptor salt bridge distances in the presence (blue) and absence (red) of heparin (C) K127 (blue) of the heparin-binding pocket (gold) forms a weak salt-bridge with D46 of the beta-trefoil core in the *apo* model (red). This interaction does not form in the heparin-bound Model 1 (blue). (D) Time series of the D46-K127 donor-acceptor salt bridge distances in the presence (blue) and absence (red) of heparin. (E) Table of intramolecular interactions unique to the heparin-binding pocket of heparin-bound hFGF1. (F) Time series of water molecule count within 3 Å of the heparin-binding pocket.

A count of the number of water molecules within 3 Å of the heparin-binding pocket provides additional evidence for the occurrence of a conformational change. Around 200 water molecules are present within 3 Å of the heparin-binding pocket throughout both heparin-bound trajectories (Figure 2F, Supplemental Figure S3E-F). During the *apo* simulation, the count of water molecules increases to 280 after 2 μs (Figure 2F), thus coinciding with the observed conformational change.

### Intramolecular interactions within the heparin-binding pocket help stabilize heparin-bound hFGF1

Thus far, we have shown that monomeric hFGF1 is destabilized in the absence of heparin and that a conformational change occurs in the heparin-binding pocket. This conformational change does not occur in the heparin-bound models, which are more stable than the *apo* model. We have identified several unique intramolecular interactions within the heparin-binding pocket that contribute to the increased stability of the heparin-bound hFGF1 models (Figure 2E, Supplemental Figure S4). Occupancies are quite similar in both Model 1 and Model 2 (Figure 2E, Supplemental Figure S4). While these interactions are also present briefly in the *apo* model, none of them meet the occupancy criteria that would allow them to be described as hydrogen bonds (Figure 2E). We propose that these intramolecular interactions may form as a consequence of intermolecular interactions between positively charged residues of the heparin-binding pocket and negatively charged residues of heparin hexasaccharide. The strength of these intramolecular interactions (occupancies between 54-97% in Model 1 and 64-98% in Model 2) might thus be a factor that prevents the conformational change observed in the *apo* model from occurring in the heparin-bound models. 65 intramolecular hydrogen bonds were observed in the *apo* model. Out of these 65, only one involves the heparin-binding pocket – L145-K142 (84%). All 65 interactions observed in the *apo* model also occur in the heparin-bound models with similar occupancies. Three of these interactions are salt-bridges – D46-R38 (90%); D53-R38 (92%); E67-K114 (60%).

Secondary structure analysis reveals that parts of the heparin-binding pocket of the *apo* model become unstructured and unravel into random coils when the conformational change occurs (Supplemental Figure S5A-B). This change in the secondary structure is then maintained for the remaining 2.8 μs of the *apo* trajectory. Residues 131 and 134-136 tend towards forming a beta strand and 3_10_ helix, respectively, prior to the conformational change. This change in the secondary structure is not observed in the heparin-bound models (Supplemental Figure S5C-D). The lack of strong intramolecular interactions in the heparin-binding pocket of the *apo* model (Figure 2E) could thus account for the observed changes in the secondary structure.

Our findings are further validated by internal RMSD analysis of the heparin-binding pocket of the *apo* (RMSD of ~4 Å) and heparin-bound (RMSD of ~0.5 Å) models (Figure 1D, Supplemental Figures S2C-D). This analysis demonstrates that the heparin-binding pockets of *apo* and heparin-bound hFGF1 have different internal conformations. Therefore, these observations confirm the role of heparin-derived intramolecular interactions in maintaining and promoting the structured nature of the heparin-binding pocket.

### Characterization of intermolecular interactions that contribute to the stabilizing effects of heparin

A visual inspection of the heparin-bound trajectories reveals that the heparin hexasaccharide in Model 1 fluctuates considerably before it eventually undergoes a 180° rotation to settle down into a more stable conformation (Supplemental Figure S6A). This transition occurs at the 1.25 μs mark and continues until the 2 μs mark (Supplemental Figure S6A). On the other hand, the heparin molecule in Model 2 does not undergo any major positional changes and attains a stable conformation very quickly (Supplemental Figure S6B). As a result of the differences in behavior and position of the heparin hexasaccharide in each model, slightly different intermolecular hydrogen bond interactions occur in each model in terms of both occupancy as well as the residues involved (Figure 3, Table 1). 6 residues in the heparin-binding pocket (R136, K132, K126, K127, R133, K142) were found to be involved in these interactions (Figure 3A-B). With the exception of R133, intermolecular hydrogen bonds involving these residues are present in the dimeric crystal structure (PDB entry: 2AXM)^32^. R133 was found to interact with heparin only in Model 1 (Figure 3A), while K142 was found to interact with heparin only in Model 2 (Figure 3B). Intermolecular interactions involving N32, N128 and Q141 are also present in the dimeric crystal structure^32^ but these residues only interact briefly (hydrogen bond occupancies < 35%) with heparin in our simulation trajectories.

**Table 1.**
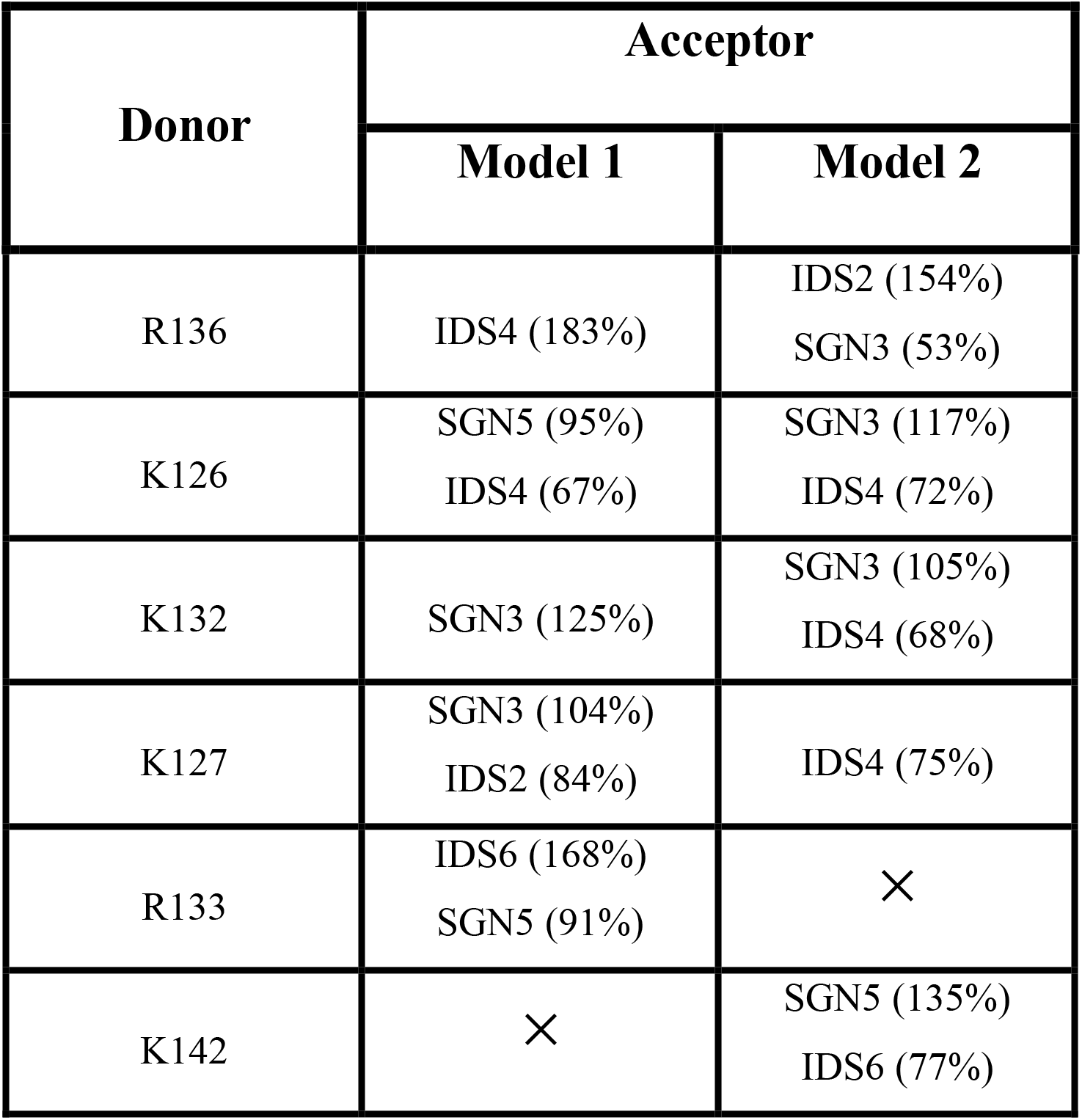
Characterization of the hFGF1-heparin intermolecular interactions in the heparin binding pocket. Table of intermolecular hydrogen-bonding interactions observed in the last microsecond of both heparin-bound trajectories. R133 interacts with heparin only in Model 1, while K142 interacts with heparin only in Model 2. Occupancies were found to be greater than 100% in several cases, indicating that more than one set of donor-acceptor atoms participate in these interactions.

| Donor | Acceptor |  |
| --- | --- | --- |
|  | Model 1 | Model 2 |
| R136 | IDS4 (183%) | IDS2 (154%)<br>SGN3 (53%) |
| K126 | SGN5 (95%)<br>IDS4 (67%) | SGN3 (117%)<br>IDS4 (72%) |
| K132 | SGN3 (125%) | SGN3 (105%)<br>IDS4 (68%) |
| K127 | SGN3 (104%)<br>IDS2 (84%) | IDS4 (75%) |
| R133 | IDS6 (168%)<br>SGN5 (91%) | × |
| K142 | × | SGN5 (135%)<br>IDS6 (77%) |

**Figure 3.**
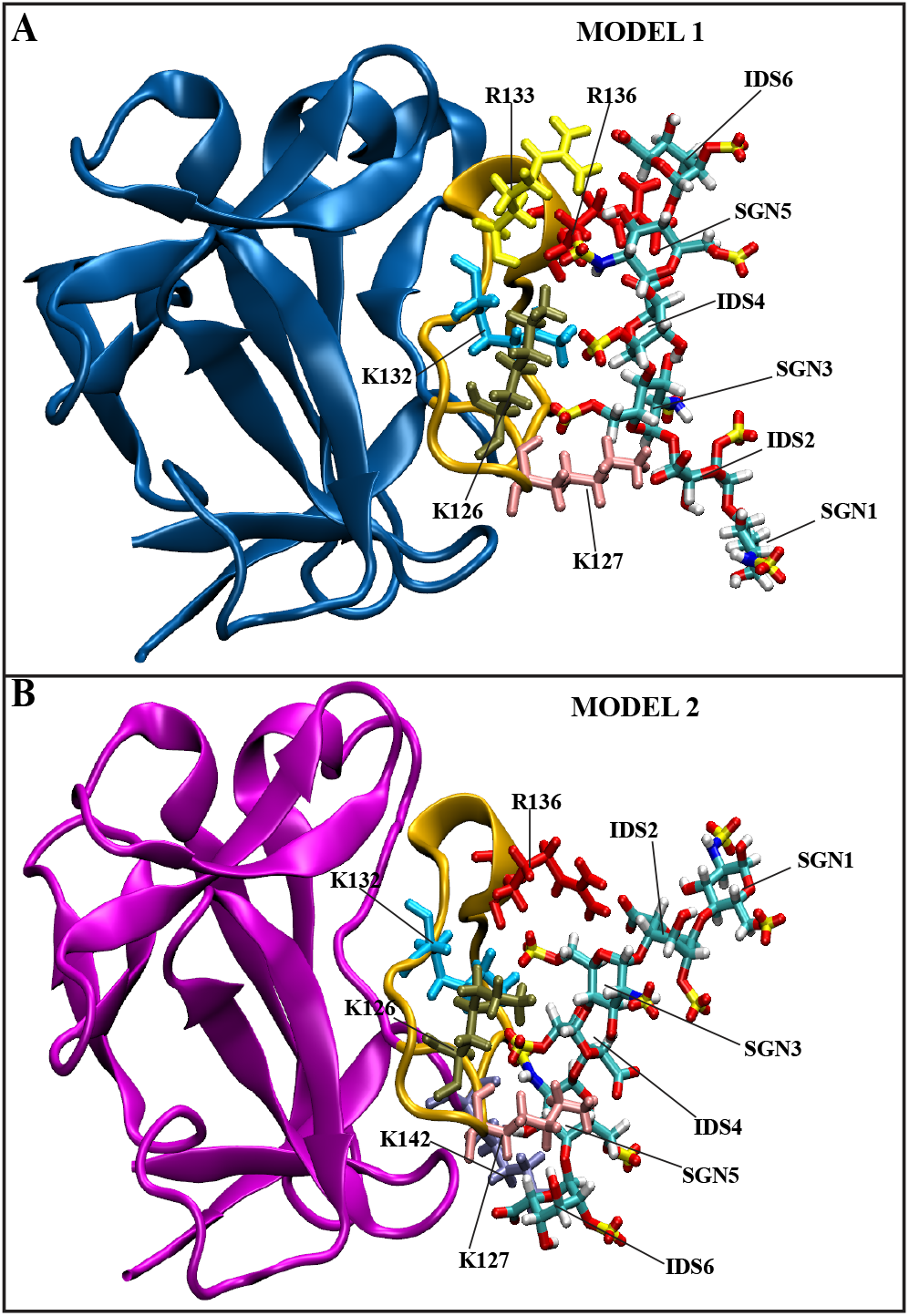
Cartoon representation of the final frames of the two heparin-bound trajectories. (A) Model 1 (blue) – heparin hexasaccharide from PDB entry 2AXM with monomeric hFGF1 from PDB entry 1RG8. (B) Model 2 (magenta) – one monomer and heparin hexasaccharide from 2AXM (dimeric). 6 residues in the heparin-binding pocket (R136, K132, K126, K127, R133, K142) were found to interact with heparin hexasaccharide.

Hydrogen bond occupancies were found to be greater than 100% in several cases (Table 1), indicating that more than one set of donor-acceptor atoms participate in these interactions. Occupancies are fairly similar for interactions involving R136 and K126 in both models (Table 1) while they are somewhat different for interactions involving residues K132 and K127 (Table 1). As discussed earlier, R133 and K142 only interact with heparin hexasaccharide in Models 1 and 2 respectively (Table 1). All 6 residues interact with both iduronic acid (IDS) and glucosamine (SGN) residues at different points during the heparin-bound simulations. The interaction of R136 with IDS2 in Model 2 (154%) is somewhat weaker than its interaction with IDS4 in Model 1 (183%) (Table 1, Figures S7A-B). The interaction of SGN3 with K132 in Model 1 (125%) and K126 in Model 2 (117%) is quite similar (Figures S7C-D).

As discussed previously, we have also identified 6 major intramolecular interactions within the heparin-binding pocket that are unique to heparin-bound hFGF1 (Figure 2E, Supplemental Figure S4). The presence of heparin ostensibly leads to the formation of these intramolecular hydrogen bonds, which consequently contribute to the stabilization of heparin-bound hFGF1.

While microsecond-level equilibrium MD simulations can be used to produce a qualitative description of protein-ligand interactions, a more accurate and efficient quantitative characterization of protein-ligand interactions can be carried out using large-scale enhanced sampling techniques. Therefore, we have calculated the absolute binding free energy for the interaction of hFGF1 with heparin hexasaccharide using a combination of SMD and BEUS simulations. The largest contributor to the absolute binding free energy was the separation of heparin hexasaccharide from hFGF1 (−18.91 kcal/mol to −16.89 kcal/mol) (Figure 4A-B, Supplemental Figure S8A). Local rotations (4.24 kcal/mol to 4.26 kcal/mol) (Figure 4B, Supplemental Figure S8B) and internal fluctuations (0.26 kcal/mol to 0.36 kcal/mol) of the heparin hexasaccharide (Figure 4B, Supplemental Figure S8D) were also considered along with the internal fluctuation contribution (0.45 kcal/mol to 0.49 kcal/mol) of hFGF1 (Figure 4B, Supplemental Figure S8C). The free energy profiles for the internal fluctuations of hFGF1 (Supplemental Figure S8C) and heparin hexasaccharide (Supplemental Figure S8D) were based on the internal RMSDs of *apo*/heparin-bound hFGF1 (Figure 1C), bound heparin and free heparin. The free energy profile based on hFGF1 internal motions clearly shows that the *apo* form of hFGF1 has two distinct conformations during the simulation (Supplemental Figure S8C). This is additional evidence for the conformational change described previously. Finally, an offset (explained in the Methods section) of ~4.44 kcal/mol needed to be added to the overall values to convert the estimated free energy to a standard binding free energy (conversion factor) (Figure 4B). The absolute binding free energy values thus ranged from −9.52 kcal/mol to −7.34 kcal/mol, given the uncertainties of different terms. In other words, our estimate for binding free energy is −8.43±1.09 kcal/mol. Binding affinity (K_d_) values calculated from the absolute binding free energy were found to be in the micromolar range with an average value of 2.3 μM and ranging from 0.12 to 4.5 μM (based on the lower and upper bounds of free energy estimates). These are in very good agreement with the K_d_ value obtained from ITC experiments (1.68±0.3 μM) (Figure 4C). The free energy value calculated from the experimental K_d_ (−7.91±0.01 kcal/mol) is also in good agreement with the computationally calculated binding free energy (−8.43±1.09 kcal/mol) (Figures 4B-C). The quantitative agreement between the computational and experimental binding affinity estimates is a great indicator of the accuracy of our extensive and multifaceted absolute binding free energy calculation method.

**Figure 4.**
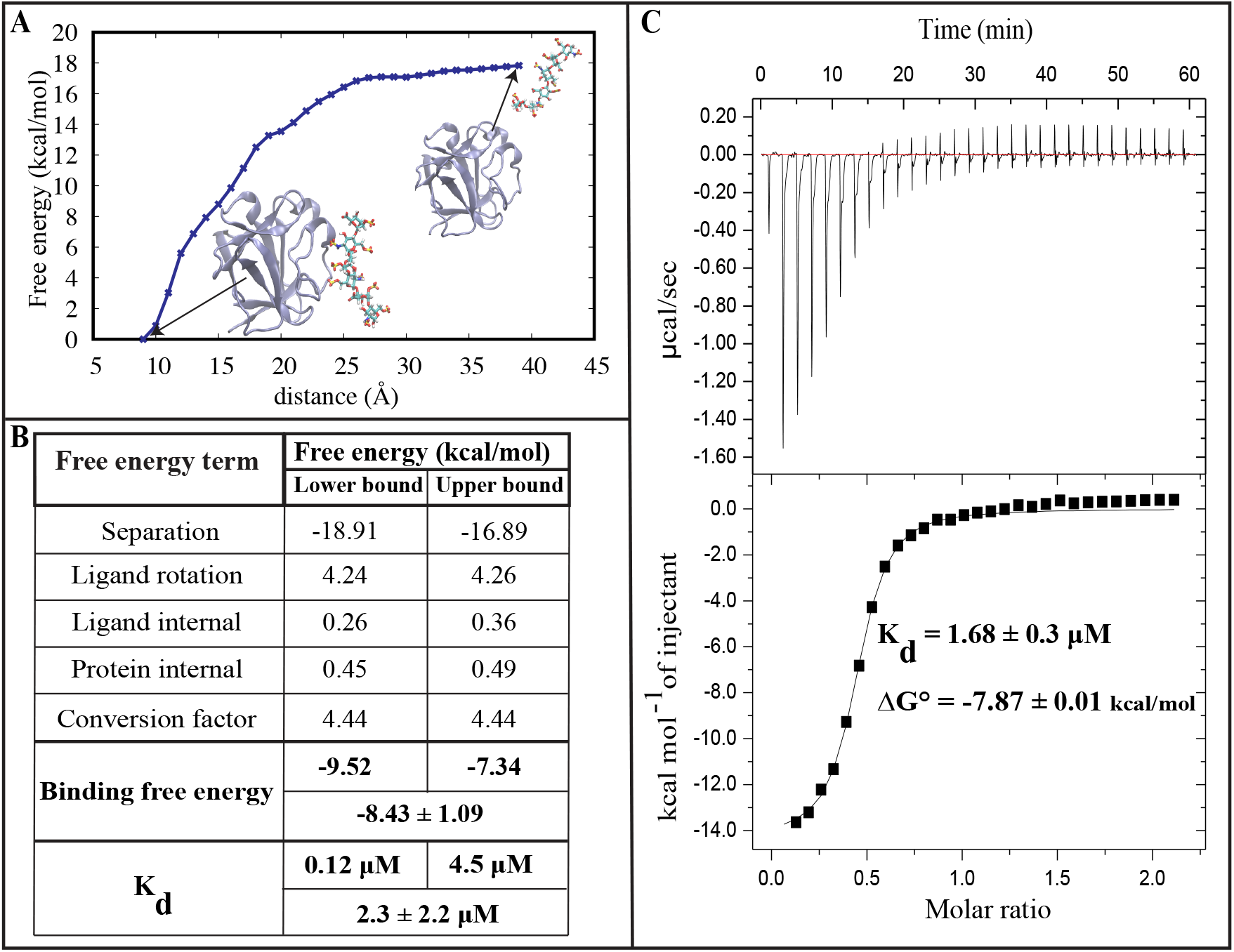
Absolute binding free energy calculations for the hFGF1-heparin interaction. (A) Free energy profile for the separation of heparin hexasaccharide from hFGF1. (B) Table of free energy terms used to calculate the absolute binding free energy. Heparin-hFGF1 separation is the largest contributor. The lower and upper bound are determined based on the uncertainty estimations in each free energy term. (C) Isothermogram representing the titration of hFGF1 with heparin hexasaccharide.

Recent computational studies have used the MM-GBSA method to calculate the binding free energy of the hFGF1-heparin interaction, with values ranging from −84.9 kcal/mol to −106.1 kcal/mol^39^. The results obtained from the MM-GBSA approach are very different from our own SMD-BEUS results, which is to be expected given that MM-GBSA ignores various contributors to the free energy, such as the entropic change that arises from the protein-ligand interactions^40,43^. Studies have shown that the binding affinity and free energy results derived from computational methods can be compared to experimental binding affinities obtained from ITC experiments^47,48^. However, for a reliable computational free energy estimate, employing purely physics-based free energy calculation methods such as those employed here has proven to be difficult. Here we show that using a careful strategy that considers all relevant free energy terms and ensures the use of powerful enhanced sampling techniques, could result in good quantitative agreements between the computational and experimental binding affinity estimates.

### Conclusions

We conclude that the conformational change observed in the heparin-binding pocket of the *apo* model is directly linked to the thermal instability displayed by unbound monomeric hFGF1. In addition to the intermolecular interactions between hFGF1 and heparin hexasaccharide, we have identified several intramolecular interactions within the heparin-binding pocket that are unique to the heparin-bound models. Thermal denaturation experiments have revealed that the T_m_ value for hFGF1 increases by approximately 20°C when bound to heparin. This suggests that the intramolecular interactions described previously play a key role in stabilizing monomeric hFGF1. We have also quantified the energetics of these interactions using ITC experiments and accurate free energy calculation methods that produced very similar results, further validating our MD based model of hFGF1-heparin interactions. Further experimental and computational research is needed to elucidate the functional relevance of these specific intramolecular interactions.

## METHODS

### Equilibrium unfolding of hFGF1 with heparin hexasaccharide

The temperature-based denaturation experiment was performed using the JASCO-1500 Circular dichroism spectrophotometer cohered with fluorescence detector. hFGF1 was diluted with 10 mM phosphate buffer containing 100 mM NaCl at pH 7.2, to get a concentration of 33μM. The experiment was performed with and without heparin. For the measurements with heparin, protein to heparin ratio of 1:10 was used. The fluorescence spectra were collected in 5 °C intervals from 25 °C to 90 °C. The fraction of denatured protein (F_d_) at each temperature was determined as F_d_ = (Y − Y_N_)/(Y_D_ − Y_N_); where, Y, Y_N_, and Y_D_ are the fluorescence signals of the 305/350 nm fluorescence ratio at the native state (25 °C), each consecutive temperature, and the denatured state (90 °C) respectively. The data set was fit using MS Excel. T_m_, the temperature at which 50 % of the protein molecules exist in the denatured state(s), was calculated from the fraction denatured protein population versus temperature graph.

### Isothermal titration calorimetry of hFGF-1 with heparin hexasaccharide

Isothermal titration calorimetry (ITC) data was obtained using MicroCal iTC 200 (Malvern Inc.). The change in heat during the biomolecular interaction was measured by titrating the heparin (loaded in the syringe) to the hFGF1 solution in the calorimetric cell. Both the protein and the heparin samples were made in the buffer containing 10 mM phosphate buffer with 100 mM NaCl at pH 7.2 and were degassed prior to loading. The protein to heparin ratio was maintained at 1:10 with the protein concentration being 100 μM and the heparin concentration being 1mM. A total of 30 injections were conducted with a constant temperature of 25 °C and stirring speed of 300 rpm. One set of sites binding model was used for the ITC binding curve^49^. The standard binding free energy ΔG° was determined from binding affinity (K_d_) via: *ΔG*° = *RT*ln(*K_d_*) at *T* = 25°C.

### All-atom equilibrium MD simulations

We have used all-atom equilibrium MD simulations to characterize the conformational dynamics of hFGF1 with and without heparin hexasaccharide. Our simulations were based on the x-ray crystal structures of the unbound hFGF1 monomer (PDB: 1RG8, resolution: 1.1 angstroms)^10^ and the dimeric complex with a heparin hexasaccharide (PDB:2AXM, resolution: 3.0 angstroms)^32^. We built three different models – monomeric *apo* hFGF1 from 1RG8; monomeric heparin-bound hFGF1 (1RG8) using the heparin hexasaccharide from the dimeric complex (2AXM) (Model 1) and monomeric heparin-bound hFGF1 from the dimeric complex (2AXM) (Model 2). Residues 12-137 in the PDB files correspond to residues 26-151 in the experimental sequence. The experiments were performed using a truncated version of hFGF1 (residues 13-154) which did not contain the unstructured 12 amino acid N-terminal segment. The unstructured N-terminal segment is not known to be involved in receptor activation or heparin binding. The heparin hexasaccharide consists of N, O6 disulfo-glucosamine and 2-O-sulfo-alpha-L-idopyranuronic acid repeats^32^.

MD simulations were performed using the NAMD 2.13^50^ simulation package with the CHARMM36 all-atom additive force field^51^. The input files for energy minimization and production were generated using CHARMM-GUI^52,53^. For heparin-bound Model 1, the heparin hexasaccharide segment from 2AXM was added to the 1RG8 structure using psfgen. The models were then solvated in a box of TIP3P waters and 0.15 M NaCl. The heparin-bound systems had approximately 23000 atoms while the *apo* system had 27,000 atoms.

Initially, we energy-minimized each system for 10,000 steps using the conjugate gradient algorithm^54^. Subsequently, we relaxed the systems using restrained MD simulations in a stepwise manner (for a total of ~1 ns) using the standard CHARMM-GUI protocol^52^. The initial relaxation was performed in an NVT ensemble while all production runs were performed in an NPT ensemble. Simulations were carried out using a 2-fs time step at 300 K using a Langevin integrator with a damping coefficient of γ = 0.5 ps^−1^. The pressure was maintained at 1 atm using the Nosé−Hoover Langevin piston method^54,55^. The smoothed cutoff distance for non-bonded interactions was set to 10–12 Å and long-range electrostatic interactions were computed with the particle mesh Ewald (PME) method^56^. The initial production run for each model lasted 15 nanoseconds, in which the conformations were collected every 2 ps. These simulations were executed on the supercomputers Razor and Trestles (University of Arkansas). After each model was equilibrated for 15 ns, the production runs were extended on the supercomputer Anton 2 (Pittsburgh Supercomputing Center) for 4.8 μs each, with a timestep of 2.5 fs. Conformations were collected every 240 picoseconds.

VMD^57^ was used to analyze the simulation trajectories. The RMSD Trajectory tool^57^ was used to calculate the RMSD and C_α_ atoms were considered for these calculations. For internal RMSD, the region of interest was aligned against its own initial configuration and RMSD was calculated with respect to this configuration. RMSF of individual residues was calculated using the C_α_ atoms by aligning the trajectory against the crystal structure. The HBond^57^ and Salt Bridge^57^ plugins were used to generate the data for hydrogen bonding and salt-bridge analysis respectively. An occupancy cutoff of 50%, a donor-acceptor distance cutoff of 4 Å and an angle cutoff of 35° were used to define hydrogen bond/salt bridge interactions. The salt bridge plugin^57^ was used to calculate the distance between the two salt bridge residues over the course of the simulation, which is the distance between the oxygen atom of the participating acidic residue and the nitrogen atom of the basic residue. The Timeline plugin^57^ was used to analyze protein secondary structure. An internal measurement method in VMD was used to count the number of water molecules within 3 Å of the heparin-binding pocket^57^.

### Multi-copy equilibrium MD Simulations for heparin hexasaccharide

The heparin hexasaccharide^32^ was also simulated in a rectangular water box without the protein. The system was setup as described previously. The final conformation after relaxation was then used as the starting conformation for 10 production runs that ran simultaneously on Blue Waters for approximately 40 ns each. The total simulation time was around 400 ns.

### Steered Molecular Dynamics (SMD)

The final conformation of the heparin-bound (Model 1) equilibrium simulation was used to generate starting conformations for the non-equilibrium pulling simulations. Two collective variables^58^ were used for two independent SMD simulations^59^: (1) distance between the heavy-atom center of mass of heparin and that of the protein and (2) the orientation-quaternion based orientation angle of heparin with respect to the protein. The distance-based SMD simulation was run for 9.5 ns, while the orientation based SMD simulation was run for 8 ns. The distance-based SMD simulation was used to pull the heparin away from the protein by approximately 30 Å (10→40 Å) with a force constant of 100 kcal/(mol.Å^2^). The orientation-based SMD simulation was used to rotate the bound heparin locally with respect to the protein (0°→73°) with a force constant of 100 kcal/(mol.*degree*^2^).

### Bias Exchange Umbrella Sampling (BEUS) simulations

Bias exchange umbrella sampling simulations^6,7,8^ were performed in order to sample as many conformations generated during the SMD simulations as possible. Selected SMD conformations were assigned to individual BEUS windows with equal spacing. The distance-based BEUS simulation ran for 10 ns with 31 replicas/windows and the orientation-based simulation ran for 10 ns with 30 replicas/windows. The force constant used was 2 kcal/(mol.Å^2^) and 0.5 kcal/(mol.*degree*^2^), respectively, for the distance- and orientation-based BEUS simulations. The rate of exchange between neighboring windows was monitored throughout the simulation. Once converged, a non-parametric weighted histogram analysis method^7,60^, which is a variation of multi-state Bennett acceptance ratio method^61^, was used to reconstruct free energy profiles.

### Absolute Binding Free Energy Calculations

Free energy calculations were performed based on data extracted from two sets of BEUS simulations described above as well as equilibrium MD simulations of heparin-bound and *apo* hFGF1. The absolute binding free energy of heparin hexasaccharide to hFGF1 was calculated using a stratification strategy, in spirit similar to that introduced by Woo and Roux^62^ and implemented recently by Fu et al.^63^ Our approach has some differences in details, particularly in the use of orientation quaternions rather than Euler angles.

We start with the calculation of the binding free energy in the simulation box. The free energy of binding involves four terms including: (1) the free energy change associated with the translation of the ligand from the binding site to the bulk (Δ*G^translation^*), the free energy change associated with the difference in rotational motion of the ligand in the binding site and in the bulk (Δ*G^ligand rotation^*), the free energy change associated with the difference in the internal flexibility of the ligand in the binding site and in the bulk (Δ*G^ligand internal^*), and the free energy change associated with the difference in the internal flexibility of the protein in the binding site and in the bulk (Δ*G^protein internal^*):

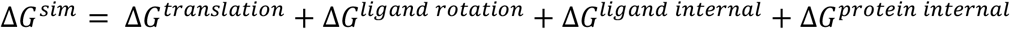

For the translational term, we use a distance-based BEUS simulation as discussed above. The BEUS-based free energy calculation estimates the separation free energy using estimated *G*(***x***), which is related to translational term via:

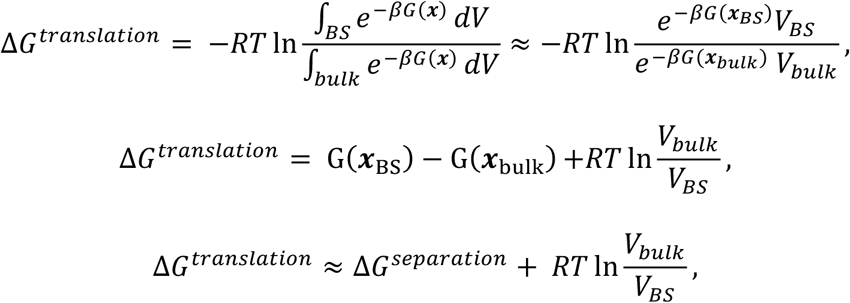

where “BS” stands for binding site. For the ligand rotational term, we use the orientation angle (Ω) within the orientation quaternion formalism, and use the orientation based BEUS to estimate *G_bound_*(Ω):

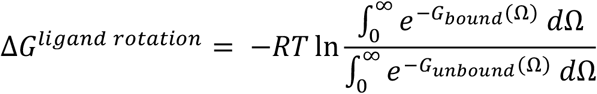

*G_unbound_*(Ω) is determined based on a freely rotating system. Finally, the internal fluctuations are estimated using RMSD time series of equilibrium simulations of free and bound ligand as well as free and bound protein:

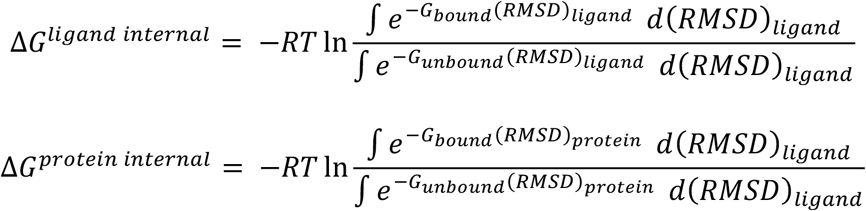

We therefore have:

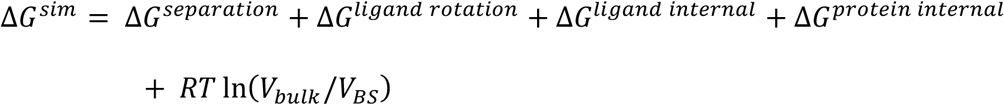

The standard binding free energy is related to the binding free energy associated with the simulation conditions via:

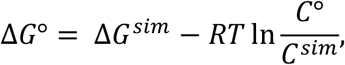

where *C*° = 1 *M* for standard condition and *C^sim^* is the ligand concentration in a simulation box of volume *V_B_* (expressed in *L*):

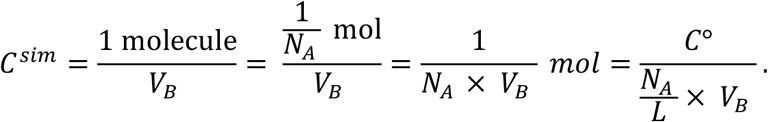

Using *N_A_* = 6.022 × 10^23^ and 1 *L* = 10^27^ Å^3^, we have:

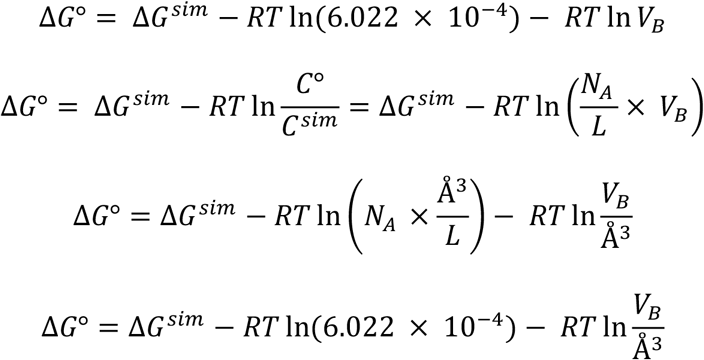

The second term on the right is evaluated at T=300 K and denoted by conversion factor:

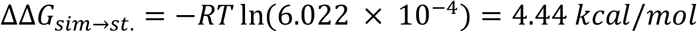

We therefore estimate the standard binding free energy to be:

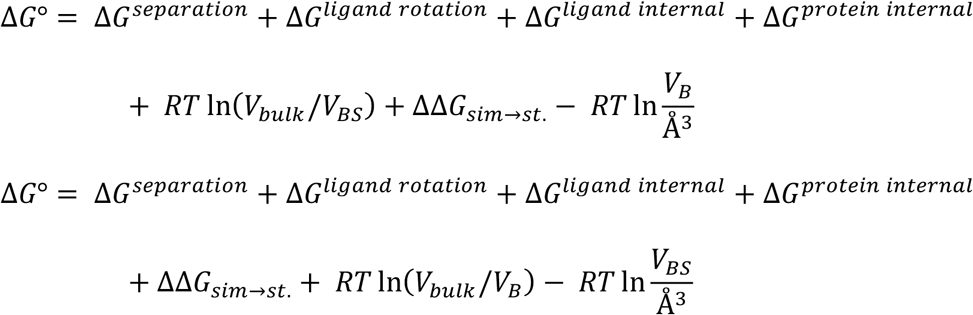

Note that the binding site volume here is the volume of the region occupied by the center of mass of the ligand when it is bound and is approximated to be around 1 Å^3^ in our simulations. We also assume *V_bulk_* ≈ *V_B_*. Therefore, we have:

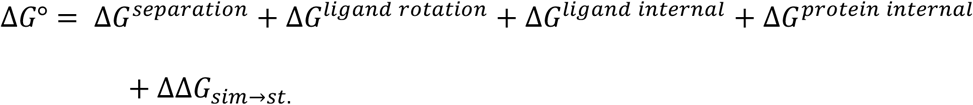

## ACKNOWLEDGEMENTS

This research is supported by National Science Foundation grant CHE 1945465 and OAC 1940188 and the Arkansas Biosciences Institute. Anton 2 computer time was provided by the Pittsburgh Supercomputing Center (PSC) through Grant R01GM116961 from the National Institutes of Health. The Anton 2 machine at PSC was generously made available by D.E. Shaw Research. This research is also part of the Blue Waters sustained-petascale computing project, which is supported by the National Science Foundation (awards OCI-0725070 and ACI-1238993) and the state of Illinois. This work also used the Extreme Science and Engineering Discovery Environment (allocation MCB150129), which is supported by National Science Foundation grant number ACI-1548562. This research is also supported by the Arkansas High Performance Computing Center, which is funded through multiple National Science Foundation grants and the Arkansas Economic Development Commission. This work is also supported by the Department of Energy (DE-FG02-01ER15161), the National Institutes of Health/National Cancer Institute (NIH/NCI) (1 RO1 CA 172631) and the NIH through the COBRE program (P30 GM103450).

## AUTHOR CONTRIBUTIONS

M.M. and T.K.S.K. designed the research. V.G.K performed the simulations and analyzed the simulation data. S.A. performed the experiments and analyzed the experimental data. V.G.K, M.M., T.K.S.K., S.A. wrote the manuscript.

## AVAILABILITY OF DATA

The molecular dynamics trajectories and the analyses generated will be shared upon request to corresponding author.

