## Supporting Information for "Mechanistic Picture for Monomeric Human Fibroblast Growth Factor 1 Stabilization by Heparin Binding"

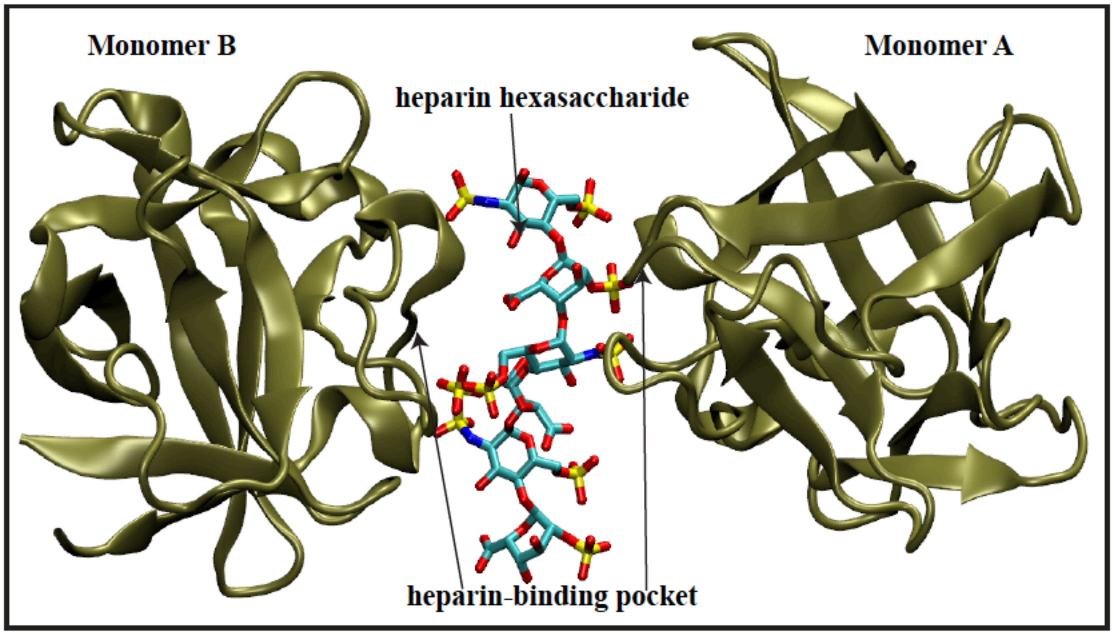


**Figure S1. Cartoon representation of the dimeric hFGF1 X-ray crystal structure with heparin hexasaccharide (PDB entry 2AXM) (related to Figures 1-4 and Table 1).**

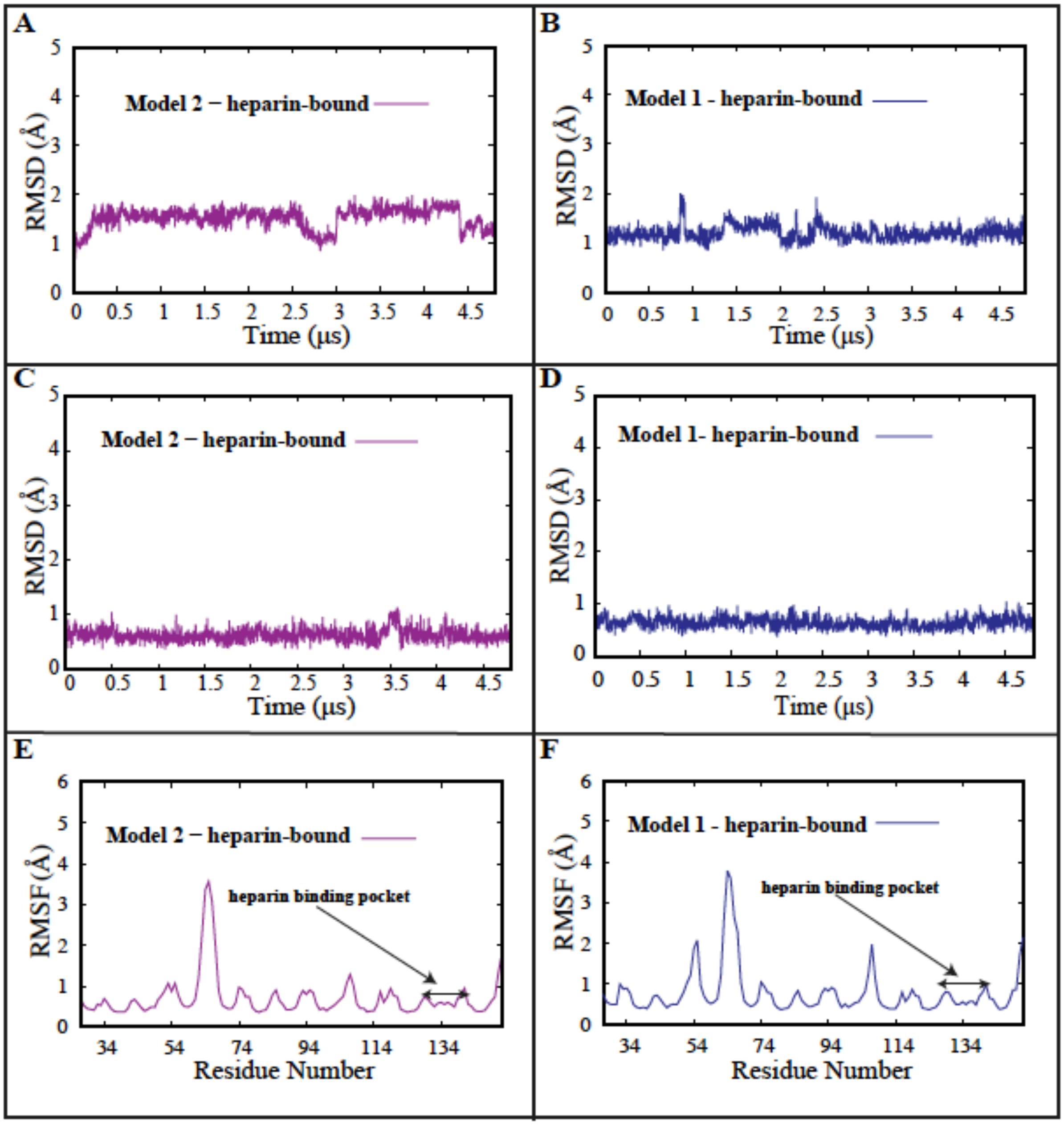


**Figure S2. Stability of heparin-bound hFGF1 assessed through RMSD and RMSF calculations (related to Figure 1).** (A) Internal RMSD time series for heparin-bound hFGF1 (Model2). (B) Internal RMSD time series for heparin-bound hFGF1 (Model1). (C) Internal RMSD time series for the heparin-binding pocket of heparin-bound hFGF1 (Model2). (D) Internal RMSD time series for the heparin-binding pocket of heparin-bound hFGF1 (Model1). (E) RMSF estimation for heparin-bound hFGF1 (Model2). (F) RMSF estimation for heparin-bound hFGF1 (Model1).

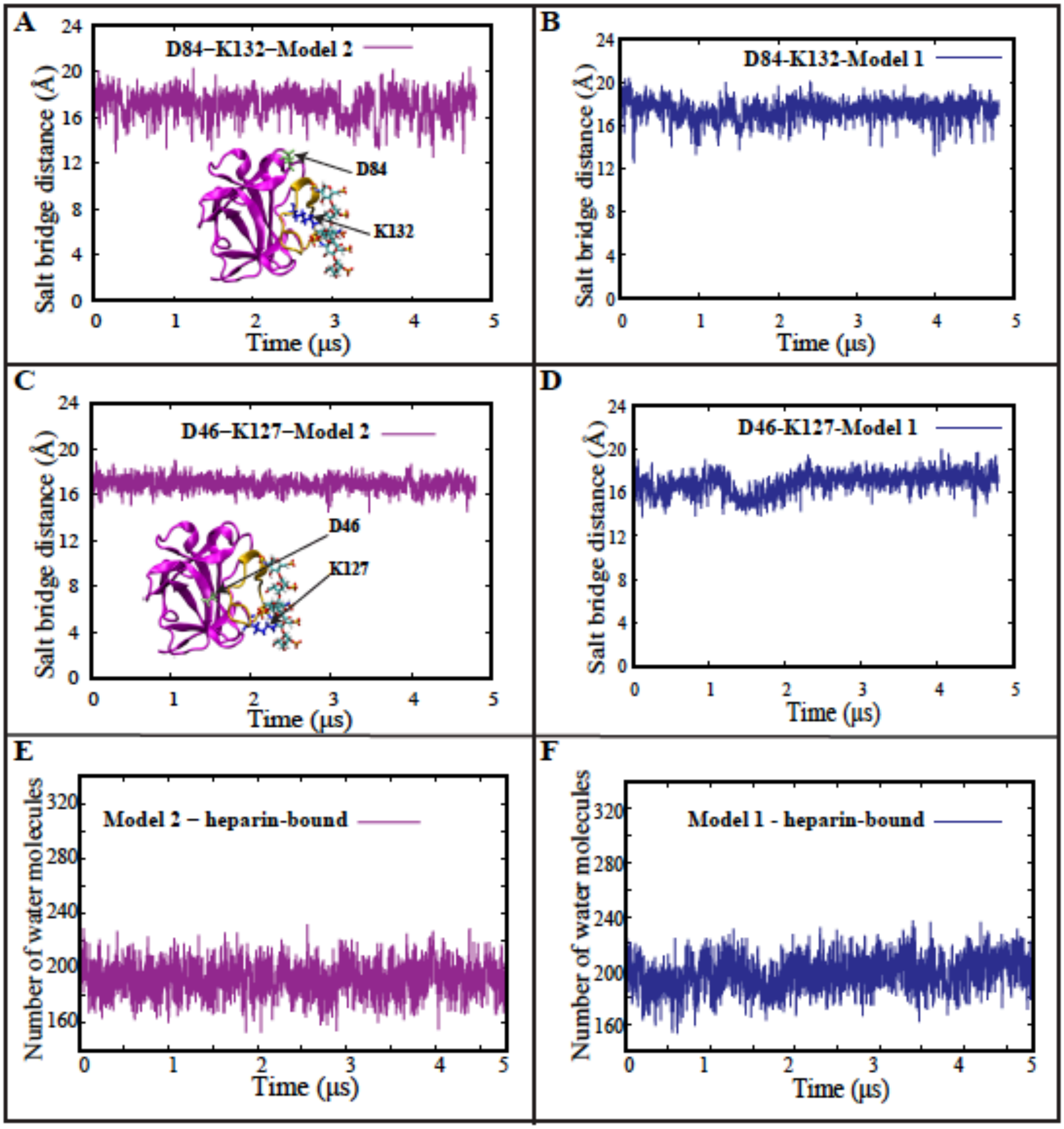


**Figure S3. Salt bridges associated with the conformational change in the apo model do not form in the heparin-bound models (related to Figure 2).** (A) Time series and cartoon representation of the D84-K132 donor-acceptor salt bridge distance for heparin-bound Model 2. (B) Time series of the D84-K132 donor-acceptor salt bridge distance for heparin-bound Model 1. (C) Time series and cartoon representation of the D46-K127 donor-acceptor salt bridge distance for heparin-bound Model 2. (D) Time series of the D46-K127 donor-acceptor salt bridge distance for heparin-bound Model 1. (E) Time series of water molecule count within 3 Å of the heparin-binding pocket for heparin-bound Model 2. (F) Time series of water molecule count within 3 Å of the heparin-binding pocket for heparin-bound Model 2.

| Donor | Acceptor | Occupancy (%) |
|---|---|---|
| K126 | S130 | 98 |
| Q141 | R136 | 90 |
| G129 | K126 | 83 |
| R136 | R133 | 74 |
| T137 | G134 | 69 |
| H138 | Q141 | 64 |

**Figure S4. Table of intramolecular interactions unique to the heparin-binding pocket of heparin-bound hFGF1 (Model 2) (Related to Figure 2E).** Hydrogen-bonding occupancies are similar in heparin-bound Model 1.

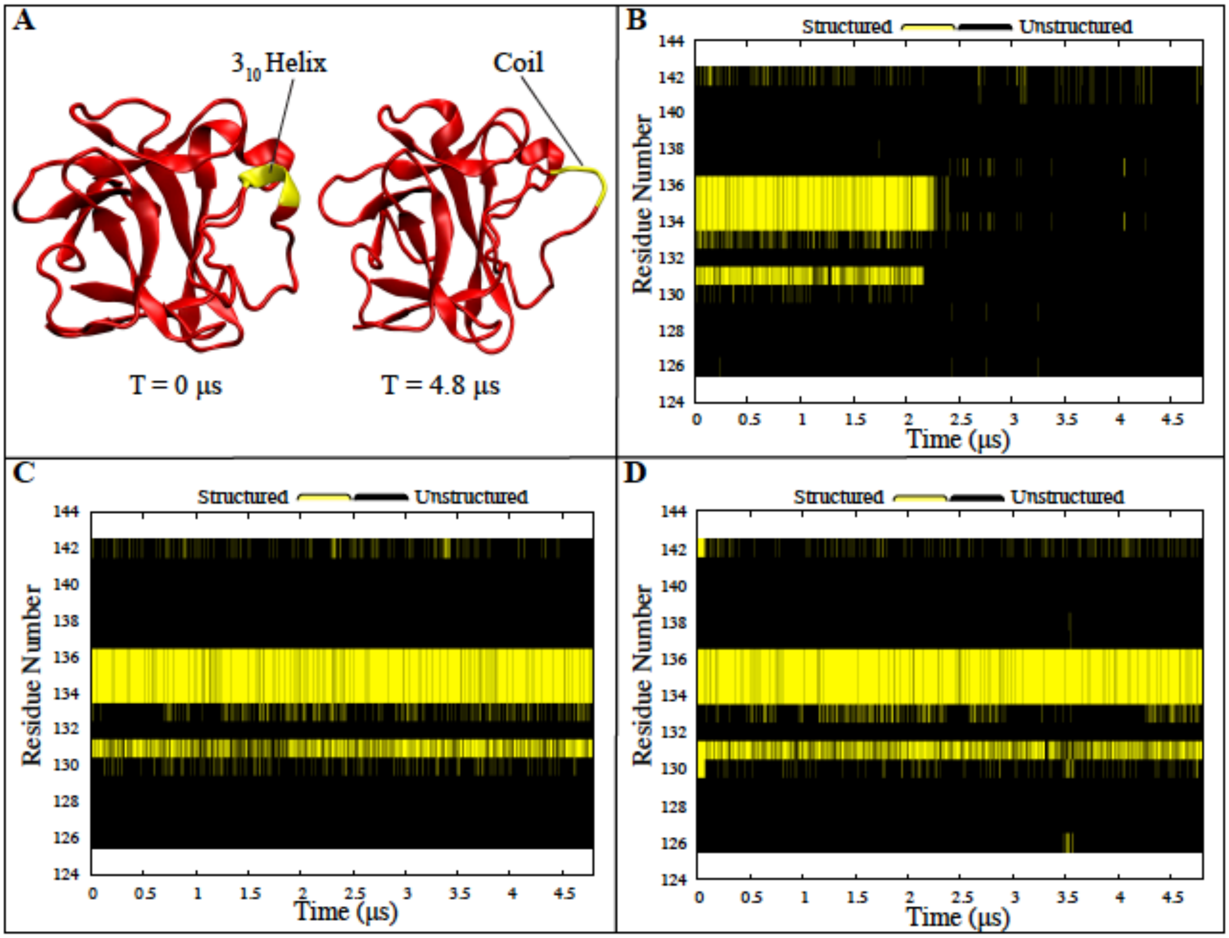


**Figure S5. The conformational change causes secondary structural changes in the apo model (related to Figure 1 and Figure 2).** (A) Cartoon representation of the secondary structural change that occurs in the heparin-binding pocket of the apo model due to the conformational change. (B) Secondary structure of the heparin-binding pocket of apo hFGF1 as a function of simulation time. Parts of the heparin-binding pocket become unstructured after 2 microseconds. (C) Secondary structure of the heparin-binding pocket of heparin-bound hFGF1 (Model 1) as a function of simulation time. (D) Secondary structure of the heparin-binding pocket of heparin-bound hFGF1 (Model 2) as a function of simulation time. The heparin-binding pocket remains structured in both heparin-bound trajectories.

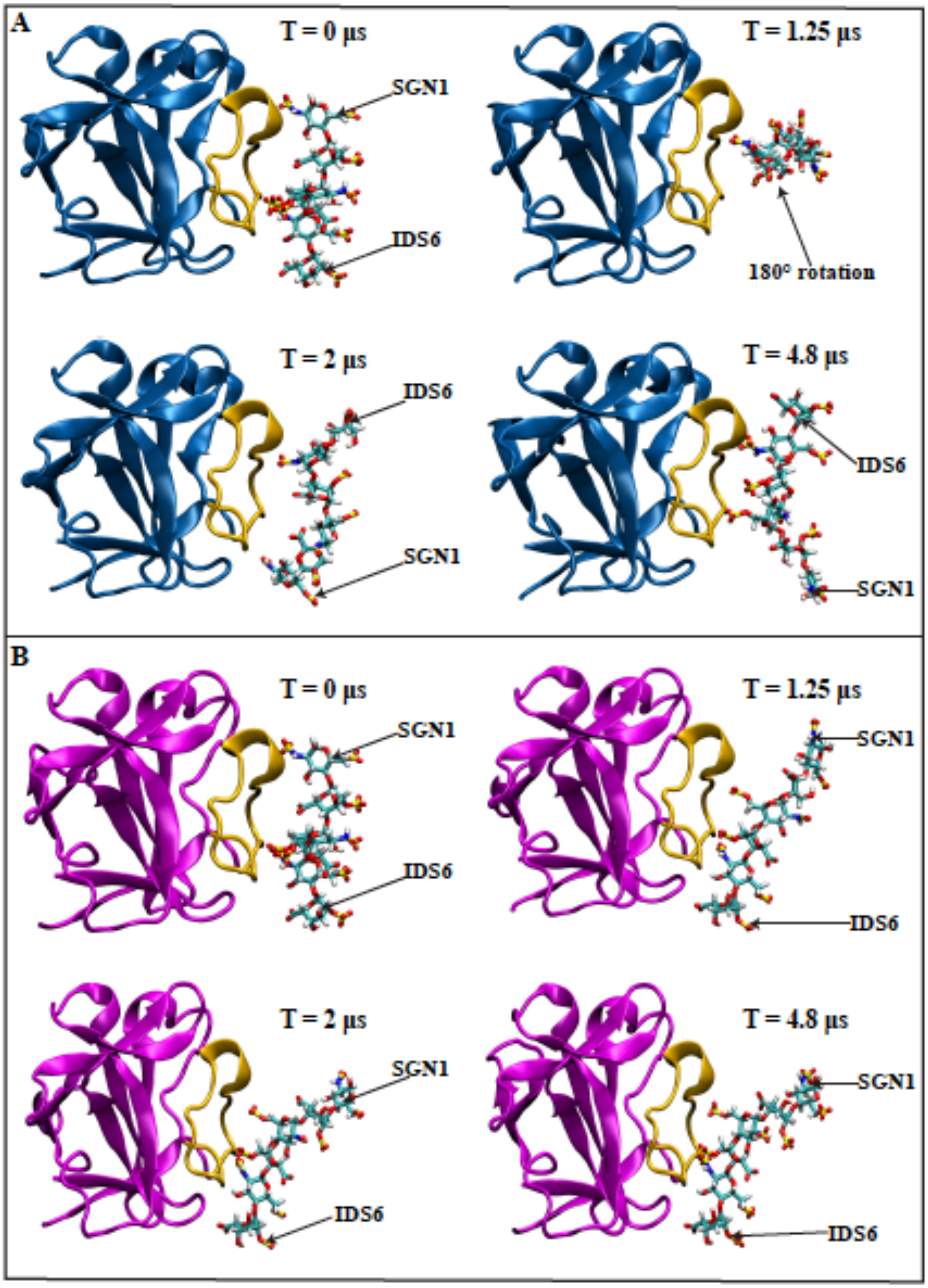


**Figure S6. Behavior of heparin hexasaccharide in the heparin-bound trajectories (related to Figure 3, Table 1 and Figure 4).** (A) The heparin hexasaccharide in Model 1 (blue) fluctuates considerably before undergoing a 180° rotation. It settles into a more stable conformation after 2 μs. (B) The heparin hexasaccharide in Model 2 does not undergo any major positional changes. Due to the differences in behavior of heparin in each model, slightly different intermolecular interactions occur in terms of both occupancy as well as the residues involved.

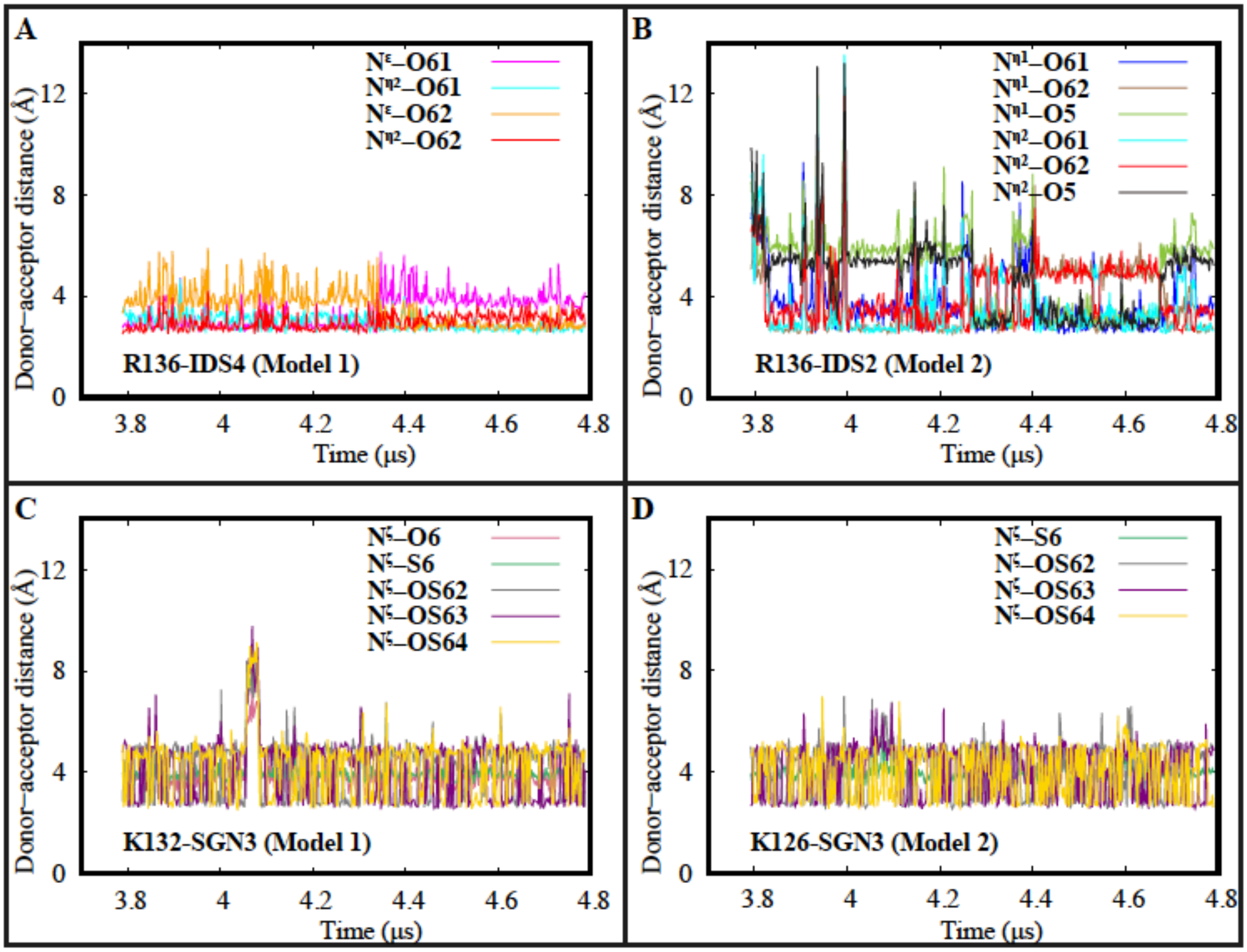


**Figure S7. Time series of hFGF1-heparin intermolecular interactions (related to Figure 3 and Table 1).** (A) Time series of hydrogen-bonding interactions between R136 and IDS4 (Model 1). (B) Time series of hydrogen-bonding interactions between R136 and IDS2 (Model 2). (C) Time series of hydrogen-bonding interactions between K132 and SGN3 (Model 1). (D) Time series of hydrogen-bonding interactions between K126 and SGN3 (Model 2).

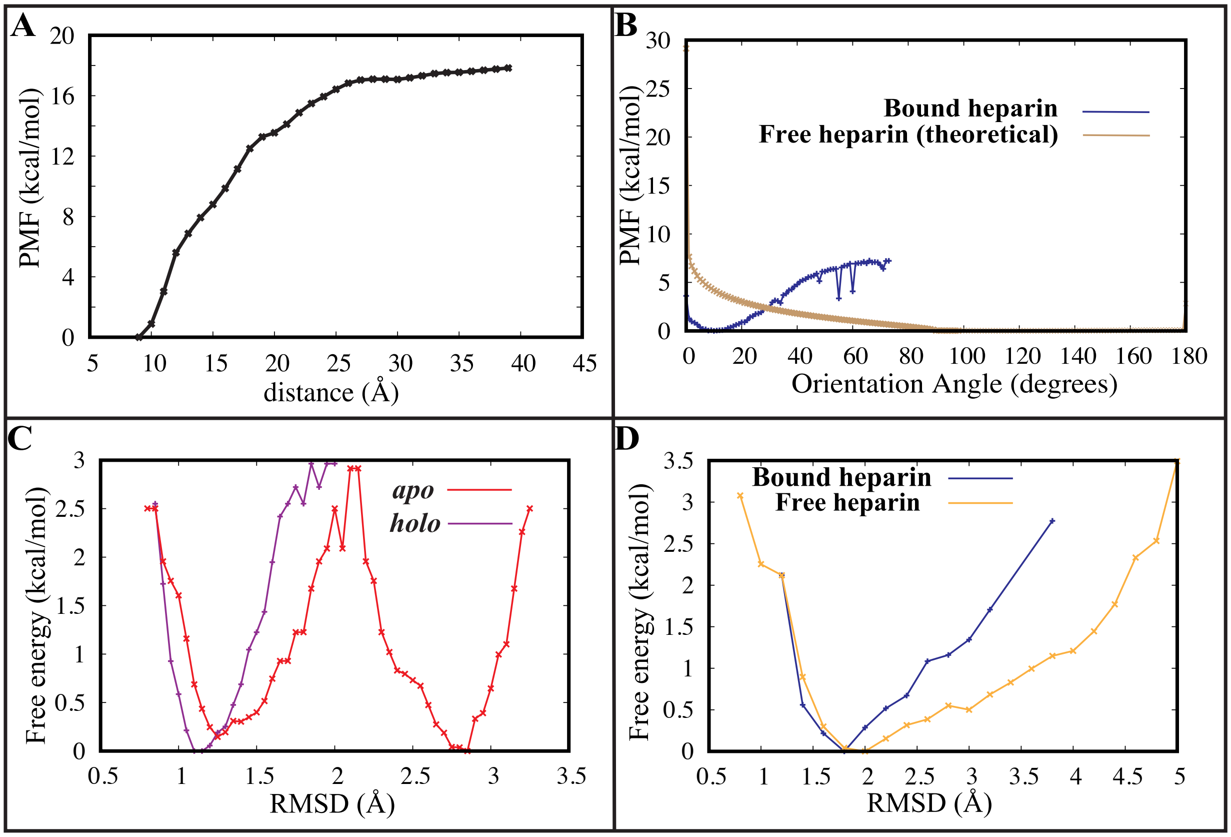


**Figure S8. Absolute binding free energy calculations for the hFGF1-heparin interaction (related to Figure 4).** (A) Potential of mean force (PMF) as a function of distance between the ligand and the heparin binding pocket (also shown in Fig. 4A). (B) Comparison of rotational PMF in bound and free forms of heparin estimated from orientation based BEUS simulations (for the bound form) and theoretically (for the free form). (C) PMF associated with internal fluctuations of apo (red) and heparin-bound (magenta) hFGF1. (D) PMF associated with internal fluctuations of free (orange) and bound (blue) heparin hexasaccharide.
